# Atomic resolution structure of serine protease proteinase K at ambient temperature

**DOI:** 10.1101/110361

**Authors:** Tetsuya Masuda, Mamoru Suzuki, Shigeyuki Inoue, Changyong Song, Takanori Nakane, Eriko Nango, Rie Tanaka, Kensuke Tono, Yasumasa Joti, Takashi Kameshima, Takaki Hatsui, Makina Yabashi, Bunzo Mikami, Osamu Nureki, Keiji Numata, So Iwata, Michihiro Sugahara

## Abstract

Atomic resolution structures (beyond 1.20 Å) at ambient temperature, which is usually hampered by the radiation damage in synchrotron X-ray crystallography (SRX), will add to our understanding of the structure-function relationships of enzymes. Serial femtosecond crystallography (SFX) has attracted surging interest by providing a route to bypass such challenges. Yet the progress on atomic resolution analysis with SFX has been rather slow. In this report, we describe the 1.20 Å resolution structure of proteinase K using 13 keV photon energy. Hydrogen atoms, water molecules, and a number of alternative side-chain conformations have been resolved. The increase in the value of *B*-factor in SFX suggests that the residues and water molecules adjacent to active sites were flexible and exhibited dynamic motions at specific substrate-recognition sites.

## Introduction

Protein structures at atomic resolution (beyond 1.20 Å) (*1*) can provide insights into the delineation of active sites residues, substrate recognition sites, and catalytic mechanisms (*2–5*). At such high resolution, the disposition and the protonation state of hydrogen atoms and a number of water molecules can be resolved, which promotes our understanding of enzyme functions directly linked to their structure (*6–13*).

High brilliance X-ray and electron sources, sensitive detectors, and cryo-cooling techniques have facilitated high-resolution structure determination. Yet the ultimate wavelength-limited resolution can result in radiation induced structure deformation. The radiation induced structural damage can be reduced by keeping the specimens at 100–110 K during the measurements. However, at low temperature, crystals may shrink, are apt to increase the sample mosaicity and induces structural deformation (*14,15*). Furthermore, in cryo-cooling X-ray analysis, cryoprotectants often block access to ligand binding sites. Since a large number of biological phenomena including enzyme reactions occur at an appropriate and optimal physiological temperature, obtaining the atomic resolution structures at ambient temperature can significantly enhance our understanding how enzymes work at such a temperature. However, the progress on high resolution structures at ambient temperature has been slow.

Here we report the 1.20 Å resolution structure of proteinase K obtained at ambient temperature. Proteinase K (EC 3.4.21.64) acts as a catalyst for hydrolysis (*16,17*) and aminolysis (*18,19*). It is a subtilisin-like serine protease possessing a catalytic triad (Ser-His-Asp) at its active site and an oxyanion hole (*20–23*). For nearly all biological organisms, serine proteases are essential in digestion, post-translational processing of secreted proteins, blood coagulation, neurotransmitters and hormones (*24*). Atomic resolution structures of these serine protease proteins at ambient temperature, to monitor room-temperature conformational ensembles that are essential for catalysis, ligand binding and allosteric regulation (*14*), can help to clarify part of their mechanistic roles biological functions. Although the atomic resolution structure of proteinase K at 0.98 Å is known at cryogenic temperature, the ambient temperature structure at a resolution greater than 1.5 Å has not been available (*20*).

We now provide the first ambient temperature, atomic resolution structure of proteinase K. Femtosecond X-ray pulses of 13□keV photon energy were used for crystallographic data collections. For serial sample loading, we employed a cellulose matrix as protein carrier with low background scattering noise. Diffraction data of ~82,000 indexed patterns obtained by SFX enable the detection of hydrogen atoms in secondary structures at ambient temperature. The proteinase K structure resolved at 1.20 Å clearly reveals flexible conformation of the residues adjacent to active sites as well as substrate binding sites and the presence of specific water molecules at ambient temperature.

## Methods

### Sample preparation

For SFX experiments, proteinase K from *Engyodontium album* (No.P2308, Sigma) was crystalized by mixing a 1:1 ratio of 40 mg ml^−1^ protein solution in 20 mM MES**–**NaOH (pH 6.5) and precipitant solution composed of 0.5 M NaNO_3_, 0.1 M CaCl_2_, 0.1 M MES**–**NaOH (pH 6.5). Microcrystals (size 8**–**12 μm) were produced by incubation for 5-10 min at 18°C. A 1.0-ml sample of crystallization solution was centrifuged at 20°C and 3,000 *g* for 3 min, and then the supernatant solution was removed. The crystals of proteinase K were suspended in 1.0 ml of the crystallization reagent. The crystal suspensions were filtered through a mesh (pore size, 30 μm). The crystal number density was adjusted to 4.9 × 10^7^ crystals/ml. The sample was stored at 18°C.

A hydroxyethyl cellulose (No.09368, Sigma) was introduced as a novel carrier matrix for SFX. After a 100-μl sample of the storage solution was centrifuged for 10 sec, a 90-μl aliquot of supernatant solution was removed. A 10-μl aliquot of the crystal solution was dispensed into 90 μl of a mixture solution of 20**–**22% (*w/v*) hydroxyethyl cellulose aqueous solution (50 μl) and the crystallization solution (40 μl) was placed on a glass slide and then mixed with a spatula. An aliquot (30 μl) of the sample was extruded into a 100-μl syringe (No.1710, Hamilton).

For synchrotron experiments, diffraction-quality crystals of proteinase K (size 100 × 100 × 200 μm) were obtained using the oil microbatch method (*25*). A crystallization drop of 1.0 μl, was created by mixing a 1:1 mixture of 40 mg ml^−1^ protein solution in 50 mM MES**–**NaOH (pH 6.5) and precipitant solution composed of 250 mM NaNO_3_, 50 mM CaCl_2_, and 50 mM MES**–**NaOH (pH 6.5) was placed in a well of a Nunc HLA crystallization plate (Nalge Nunc International) which was then covered with 20 μl of paraffin oil.

### SFX data collection

We carried out the experiments using femtosecond X-ray pulses from the SPring-8 Angstrom Compact Free Electron Laser (SACLA). The X-ray wavelength was kept at 0.95 Å (13 keV) with a pulse energy of ~200 μJ. Each X-ray pulse delivers ~7 × 10^10^ photons within a 10 fs duration (FWHM). Data were collected using X-ray beams of 1.5 × 1.5 μm^2^ focused by Kirkpatrick–Baez mirrors (*26*). The crystals in the matrix were serially loaded using a syringe injector (*27,28*) installed in a helium-ambiance diffraction chamber. The experiment was carried out using the Diverse Application Platform for Hard X-ray Diffraction in SACLA (DAPHNIS) (*29*) at BL3 (*30*). The microcrystals embedded in the matrix were kept at a temperature of approximately 20 or 4°C in a sample injector. The sample chamber was kept at a temperature of ~25°C and humidity greater than 80%. Diffraction patterns were collected using a custom-built 4 M pixel detector with multi-port CCD sensors (*31*). The cellulose matrix with randomly oriented crystals was extruded through an injector nozzle with an inner diameter of 110 μm. The sample flow rate was 0.34**–**0.46 μl min^−1^. Data collection was guided by real-time analysis in the SACLA data processing pipeline (*32*).

Diffraction patterns were filtered and converted by Cheetah (*33*) adapted (*32*) for the SACLA data acquisition system (*34*). Each pattern with more than 20 spots was accepted as a hit, and indexed and integrated using CrystFEL version 0.6.2 (*35*). The detector metrology was refined by geoptimiser (*36*). Diffraction peak positions were determined using the built-in Zaefferer algorithm (threshold 400, minimum gradient 500,000, minimum signal-to-noise 1) and passed on to DirAx 1.16 for indexing (*37*). Images with estimated resolution worse than 1.9 Å were rejected. Measured diffraction intensities were merged by *process_hkl* in the CrystFEL. No sigma cutoff or saturation cutoff were applied in the Monte-Carlo integration (**table S1**).

### SRX data collection

A proteinase K crystal was placed in a cold nitrogen gas stream after a brief soaking in a cryoprotectant solution: 30%(*w/v*) glycerol in the precipitant solution. X-ray diffraction images were collected using an RAXIS-V area detector (Rigaku, Tokyo, Japan) with synchrotron radiation in the wavelength of 0.80 Å at the BL-26B1 station of SPring-8 (Hyogo, Japan) (*38*). The data obtained were processed, merged, and scaled using the HKL2000 program package (*39*).

### Structure determination for SFX and SRX data

Each structure for SFX and SRX was determined by difference Fourier synthesis using a search model (PDB code: 4b5l). Manual model revision was performed using the Coot (*40*) program. The program PHENIX (*41*) was used for structure refinement. Water molecules were incorporated where the difference in density exceeded 3.0σ above the mean and the 2m*F_o_***–**D*F_c_* map showed a density of more than 1.0σ. All reflections were included with no σ cutoff; 5% of the data were randomly selected and omitted during refinement for cross validation by means of the free *R*-factor. The occupancy of the major conformation was refined first, and then the minor conformation was assigned and refined based on its m*F_o_***–**D*F_c_* map. Anisotropic *B*-factor refinement was performed, and finally hydrogen atoms were automatically added to the models. Hydrogen atoms were included in the protein and glycerol atoms but not in the nitrate/solvent atoms. The hydrogen omit maps were created by deleting all the hydrogen atoms from the model and refined through PHENIX.

The quality of the final model was assessed using PROCHECK (*42*) and RAMPAGE (*43*). The CCP4 package was used for the manipulation of data and coordinates (*44*). The electron density maps and structural images were generated using PyMOL (*45*). The comparisons of water molecules were performed by WATERTIDY implemented in CCP4. The coordinates and observed intensities of proteinase K have been deposited in the PDBj (accession code 5kxu for SFX and 5kxv for SRX). The total absorbed dose was calculated by RADDOSE (*46*). Details of data collection and refinement statistics are shown in **Table 1**.

**Table 1.**
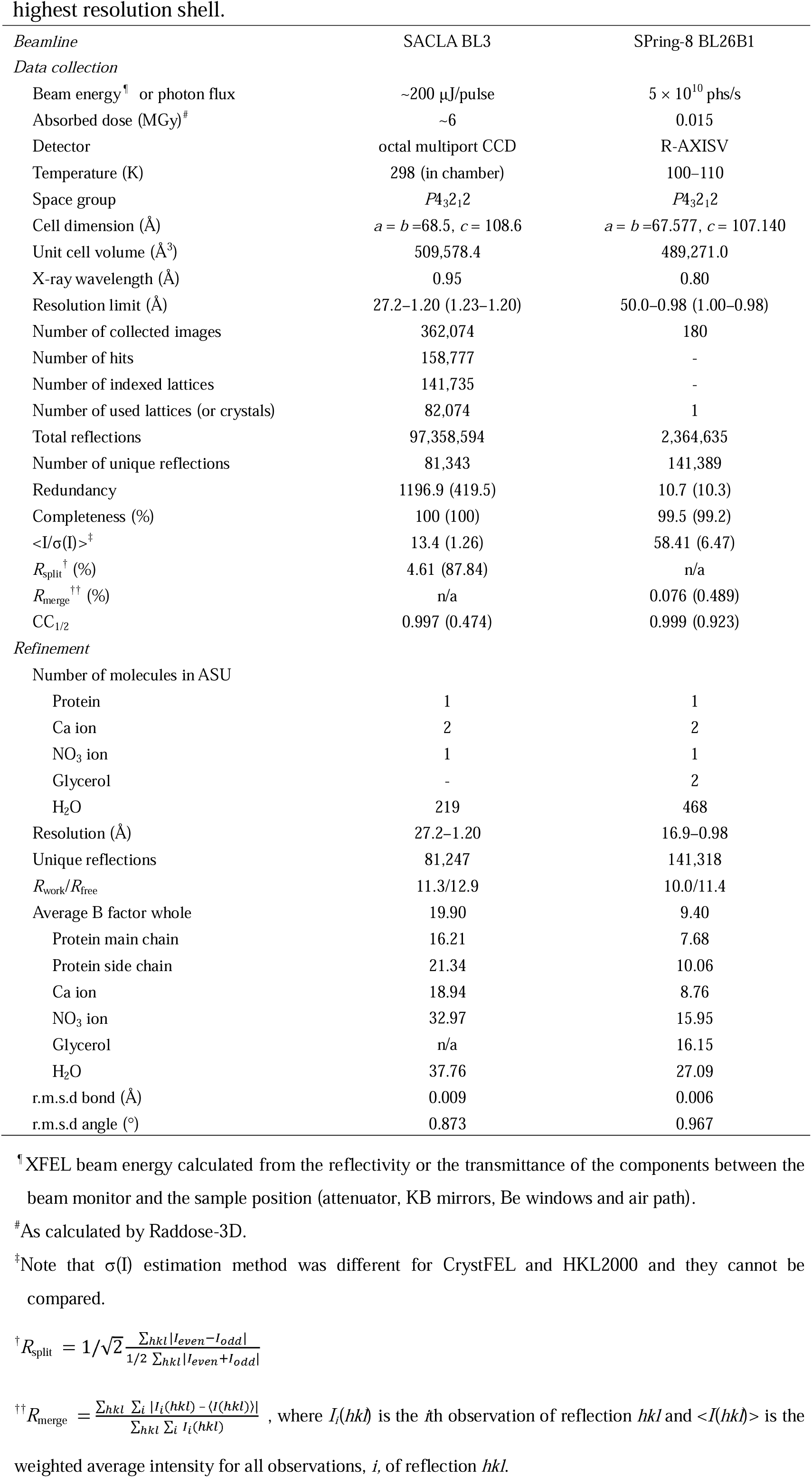
| Data collection and refinement statistics. Values in parenthesis are for the highest resolution shell.

### Results and Discussion

#### Hydrogen atoms in the secondary structures detected at room-temperature

Using ~82,000 indexed patterns at 13 keV photon energy (**fig. S1**), the room-temperature structure of proteinase K was determined to a resolution of 1.20 Å by SFX (**Fig. 1**). To determine the ‘high resolution limit’, we performed paired refinement tests based on the theory that proper use of weaker and noisier, but higher-resolution data, can provide better models (*47,48*). The inclusion of noisy high-resolution data (**fig. S2a,b**) produces a better analysis of SFX data (*49,50*). The result of paired refinement tests showed that including higher-resolution data provided a better model than rejecting it (**fig. S2c**). We therefore chose to use data up to 1.20 Å resolution. We noticed that intensity fall-off at higher resolution shells was quicker than the prediction from the Wilson plot. This was possibly caused by two factors. First, the still-Lorentz factor is not modeled by CrystFEL. Second, variations in *B* factors are not scaled by Monte Carlo integration (*50*).

**Fig. 1.**
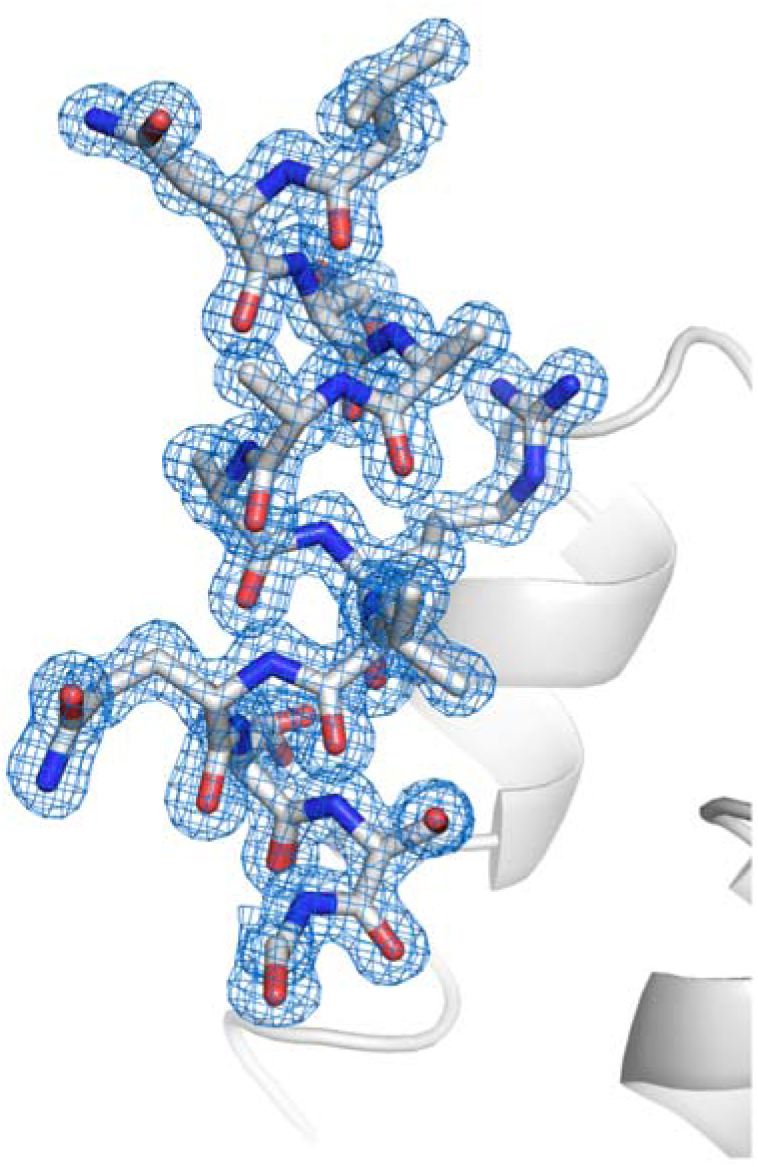
**A close-up view of the proteinase K structure with a 2m*F*_o_–D*F*_c_ electron-density map contoured at the 1.5_σ_ level.**

The structure was refined anisotropically and the final *R*/*R*_free_ factors converged to 11.3/12.9% (**Table 1**). Among 2,185 hydrogen atoms, about 250 atoms could be detected in the hydrogen omit map. Hydrogen atoms forming hydrogen bonds in the α-helix, β-sheet and turn have been successfully assigned (**Fig. 2a–c**). The electron densities of hydrogen atoms in helix (n) N-H⋯O=C (n-4) were refined for helices α3, α4 (**Fig. 2a**), and α5. Similarly, hydrogen atoms in the β-sheet were located in the central pleated β-sheet between βII1 to βII3 (**Fig. 2b**). The electron densities for hydrogen atoms in the turns (n) N-H⋯O=C (n-3) and (n) N-H⋯O=C (n-4) were detected in type I β-turn, and designated t17 consisting of Glu196–Leu199 **(Fig. 2c)**.

**Fig. 2.**
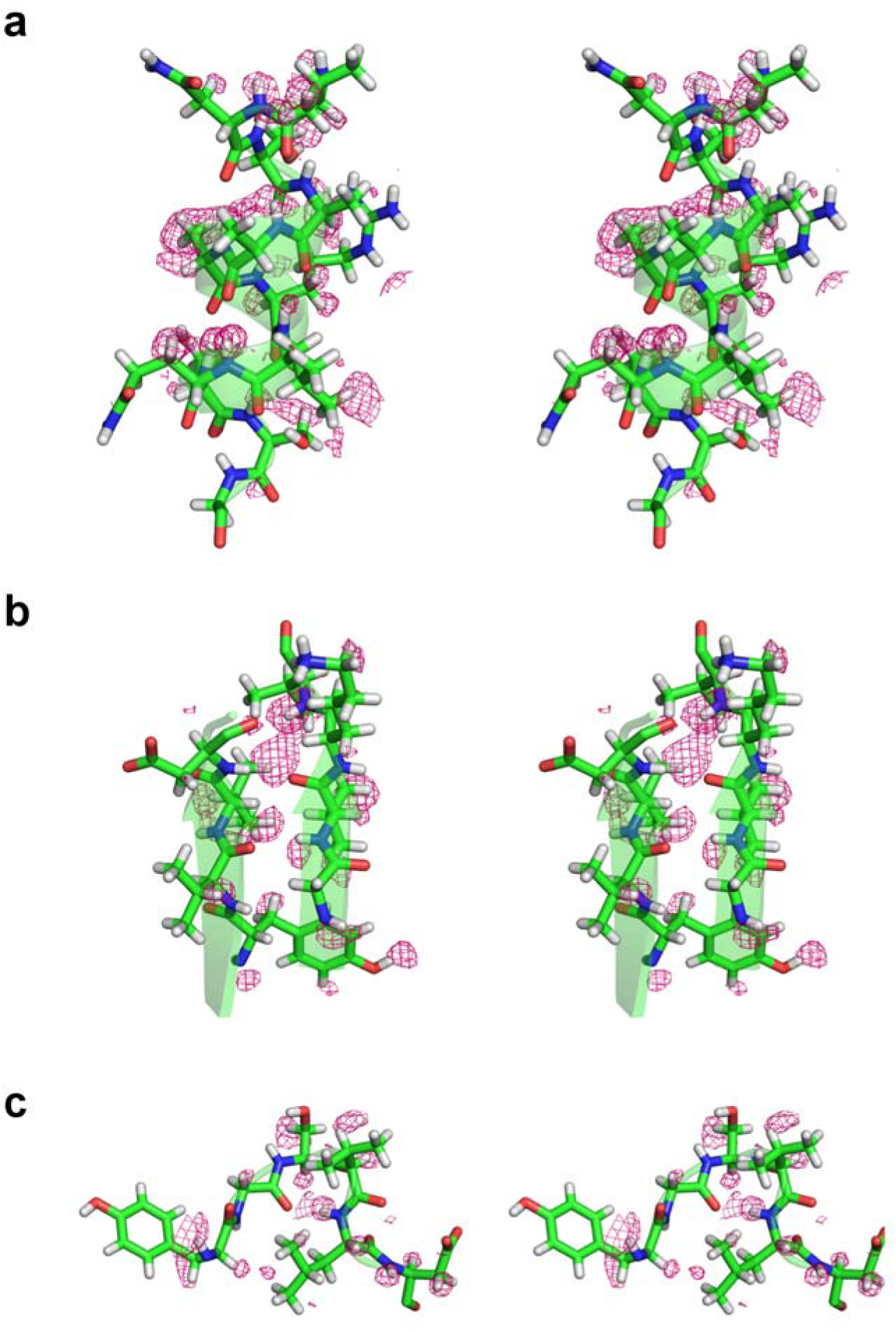
Stereo views of hydrogen atoms in secondary elements in SFX structure. Typical hydrogen atoms are assigned in SFX structure, in (**a**) α-helix, (**b**) β-sheet, and (**c**) turn. The σA-weighted m*F_o_***–**D*F_c_* maps omitting hydrogen atoms contoured at 2.0σ are shown in pink.

To compare the structures obtained via SFX and synchrotron radiation crystallography (SRX), we also collected cryo-cooled high resolution data using synchrotron X-rays. We determined the structure of proteinase K at the resolution of 0.98 Å by SRX (**Table 1**). In SFX structure, hydrogen atoms were visible in the α-helix, similar to the SRX structure (**table S2**). Substantial differences between SFX and SRX structures were also found in disulfide bonds as well as the Ca binding site, which are responsible for the stability and activity of proteinase K (**fig. S3**). These results indicate that the SFX approach produces minimal radiation damage at ambient temperature and can determine the atomic positions with high accuracy. To overcome the radiation damage problem, helical and grid data collection using synchrotron sources were developed (*51,52*). Recently, synchrotron-based serial crystallography data collection at room temperature has also been demonstrated (*53–55*). This development enables structure determination with minimal radiation from cryo- or room-temperature crystals. However, XFEL with ultrafast X-ray pulses allows data collection from highly radiation-sensitive proteins (*56–58*).

#### Appreciable extent of flexibility in the substrate recognition site revealed by SFX

SFX data have enabled accurate modeling of alternative conformations in side-chains. Altogether, 22 of the 279 residues have been modeled in two conformations and four serine residues have been modeled in three conformations. The most prominent feature of the alternative conformations was the serine residues, which represented more than 30% (eleven residues) among the alternative conformations. Seven asparagine, three aspartic acids, two glutamine and glutamic acids residues were also had alternative conformations. These alternative conformations were randomly distributed in a molecule. The disordered of Asn161 might be related to the position of the oxyanion water (W46) to initiate the substrate catalysis for Ser224, which is a member of the catalytic triad (**Fig. 3**). The oxygen atom (Oγ) of Ser224 is also hydrogen bonded to a nitrogen atom (Nε2) of His69. In the SFX structure, the distance was 2.99 Å and is similar to the SRX structure (2.98 Å). Three water molecules (W46, W123 and W132) participate in hydrogen bonded to catalytic residues of His69 and Ser224. An electron density map of these three water molecules are visible when contoured at 1.5σ (**Fig. 3a**), suggesting that the three water molecules in the SFX structure are stabilized by two components from His69, Ser224 and Asn161. In contrast, one of the three water molecules (W132) was not involved in the hydrogen bond formation in the cryo-SRX structure. The configurations and distances of these water molecules were investigated in detail and the SFX and cryo-SRX data was compared. The distance between the oxygen atom (Oγ) Ser224 and oxyanion water (W46) and W123 in SFX was 2.93 Å and 3.08 Å, respectively (**Fig. 3b**). These were slightly longer than the cryo-SRX data. Whereas one of the three waters in SFX (W132) had behaved quite differently when compared to the cryo-SRX data. Although in SFX, the distance between W132 and the Ser224 was 2.93 Å, the distance between Ser224 and the nearest the water in the cryo-SRX was 3.87 Å, which is considerably longer than the SFX data. This water (W132) is also stabilized by the Asn161 which possesses alternative side-chain conformations. In SFX, the distance was 3.26 Å which is longer than the SRX data (2.59 Å). Thus, it appears that the configurations of waters in the oxyanion hole are different between ambient temperature and cryo-cooled temperature, and our atomic resolution analysis enabled us to assign the presence of three water atoms how interact with one of the catalytic residues of Ser224 at ambient temperature.

**Fig. 3.**
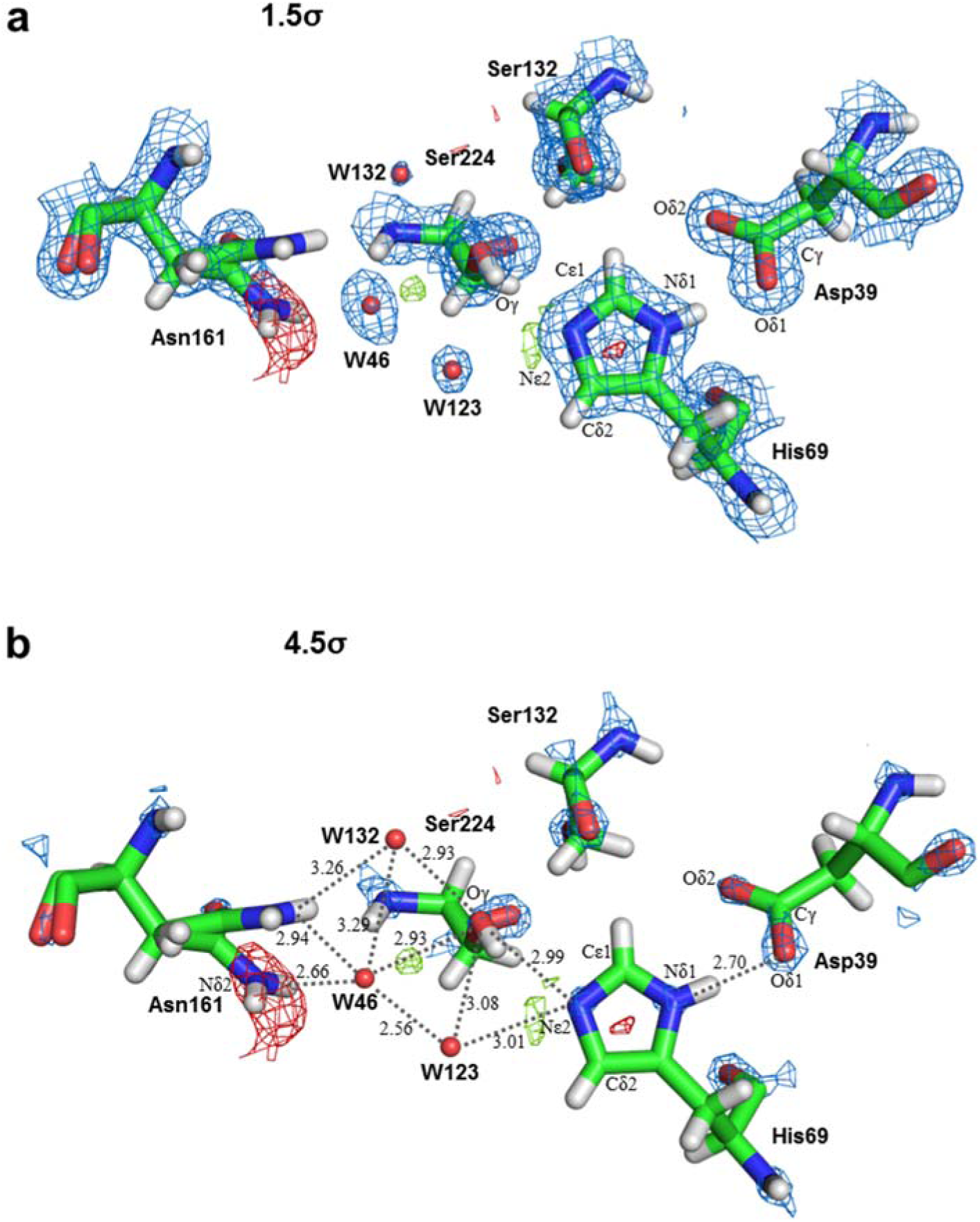
Electron density maps around catalytic triad and oxyanion hole in SFX structure. The σA-weighted 2m*F_o_***–**D*F_c_* maps contoured at (**a**) 1.5σ, (**b**) 4.5σ, are shown in blue meshes. The σA-weighted m*F_o_***–**D*F_c_* maps contoured at 2.5σ and −2.5σ are shown in green and red meshes, respectively.

Water molecules play significant roles in the structural integrity of proteins, and are buried at varying depths in the interior of molecules. In proteinase K, water clusters are involved in stabilization and assist with connecting surface segments through hydrogen bonding (*20*). As mentioned above, the locations and the configurations of waters molecules in SFX are relatively different when compared to the cryo-SRX structure; however, it was still unknown whether these differences could also be detected when compared against the non-cryo cooling SRX structure. To clarify these problems, our SFX structure was compared to two non-cryo cooled SRX structures [PDB: 2prk (*59*) and 4b5l (Jakoncic, J. *et al*. 2012): Dose ~0.04 MGy]. The distance between the oxygen atom (Oγ) Ser224 and oxyanion water for 2prk and 4b5l was 2.86 Å and 2.87Å, respectively, and the distance between the nitrogen atom (Nδ2) Asn161 and oxyanion water for 2prk and 4b5l was 2.73 Å and 2.65Å, respectively. These values are similar when compared to the SFX structure [2.93 Å for (Oγ) Ser224 and, 2.66 Å for (Nδ2) Asn161A] (**Fig. 3b**), suggesting that the location of the oxyanion water was conserved in the data from non-cryo conditions. In the SFX structure, the oxyanion water molecule (W46) also hydrogen bonded to the water molecules, W123 and W132, at distances of 2.56 Å and 3.29 Å, respectively. W123 was also engaged in hydrogen bonding to (Oγ) Ser224 (3.08 Å) and (N□2) His69 (3.01 Å). W132 was also hydrogen bonded to (Oγ) Ser224 (2.93 Å) and (Nδ2) Asn161B (3.26 Å).

In the SRX structure (PDB: 2prk), oxyanion water molecule also forms hydrogen bonding to another water molecule (2.78 Å) (*20*), but this water molecule does not seem to be engaged in the hydrogen bonding to (Oγ) Ser224 (3.74 Å) or (N□2) His69 (3.46 Å). Furthermore, no water molecule such as W132 in SFX was present around Ser224. As to the SRX structure (PDB: 4b5l), oxyanion water molecules also form hydrogen bonding to another water molecule (2.26 Å) (*20*), which is in turn also hydrogen bonded to (Oγ) Ser224 (2.75 Å) and (N□2) His69 (2.83 Å), similar to the SFX structure. Another water molecule located in a similar position as W132 in SFX was also hydrogen bonded to (Oγ) Ser224 (3.19 Å), but not hydrogen bonded to (Nδ2) Asn161B (3.72 Å).

Thus locations of water molecules in oxyanion hole, except oxyanion water, are quite different even under non-cryo conditions, we next investigated the differences in the distribution of each water molecule. In order to extract the differences in locations of water molecules between SFX and SRX, we selected water molecules which are located within a distance of 1.00 Å between SFX and SRX, and then omitted them. As shown in **Fig. 4a**,**b**, a number of SFX-specific water molecules (magenta) are distributed in molecules and notably SRX-specific water molecules (cyan) are also observed. It has been suggested that water molecules in substrate recognition regions as well as prime recognition regions play significant roles in forming water clusters. However, a few water molecules were detected in previous non-cryo SRX structures in the absence of a substrate (*60,61*). However our SFX atomic resolution structure revealed, for the first time, several water molecule candidates in these regions.

**Fig. 4.**
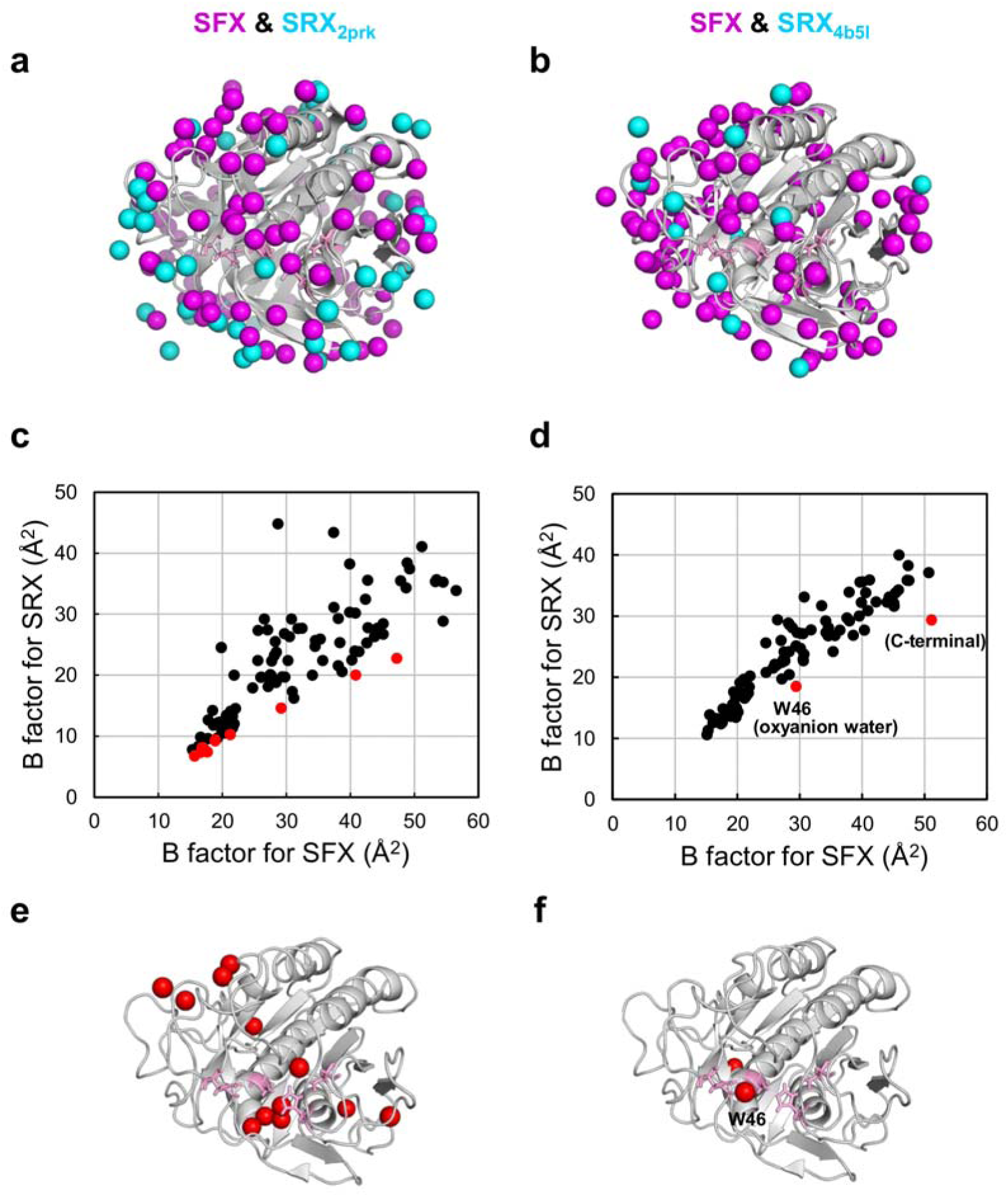
Comparison of water molecules in SFX and SRX structures. (**a,b**) Water molecules specific for SFX and two room temperature SRX structures [PDB codes: 2prk (**a**) and 4b5l (**b**)] are indicated in magenta and cyan, respectively, as sphere models. (**c,d**) All water molecules found in both SFX and SRX structures [(**c**) 2prk and (**d**) 4b5l] which are located in the positions within the distance of 1.00 Å are plotted based on their *B* factor. Striking increases of *B* factor of water molecules are indicated in red. (**e,f**) Substantial higher *B* factor of water molecules [for (**e**) 2prk and (**f**) 4b5l] are indicated in red as sphere model. Catalytic residues are show in pink as stick models.

As to water molecules found in both SFX and SRX structures located in positions within a distance of 1.00 Å were further analyzed based on their *B* factor. The average *B* factor for water molecules in the SFX structure was 37.76 Å^2^, and was larger than the SRX structure of 2prk (27.25 Å^2^) and 4b5l (25.80 Å^2^); suggesting that water moleculues in the SFX structure are highly flexible structures. Moreover, a striking increases of *B* factor was found in the interior part of molecule (2prk, **Fig. 4e**) or in two water molecules (4b5l, **Fig. 4f**) in the SFX structure when compared to water molecules in the SRX structures (**Fig 4c**,**d** shown in red). As mentioned, the location of oxyanion water is strictly conserved among cryo-SRX as well as non-cryo-SFX and SRX-structures, but it is noteworthy that the *B* factor of oxyanion water in SFX (29.36 Å^2^) was approximately 1.5 times larger than that in SRX (19.70 Å^2^ for 2prk, 18.47 Å^2^ for 4b5l), SFX suggests that oxyanion water are reliable to initiate the catalytic reaction as soon as the substrate enters the oxyanion hole. The biological relevance of the configurations and behaviors of side-chain atoms as well as the waters in the oxyanion hole might be the result the number of crystals used and their random behavior in the SFX data.

#### Dynamic motion at substrate-binding sites revealed by SFX

In order to assess the differences in structural features between SFX and SRX more clearly, both structures were superposed and r.m.s.d. was investigated (**fig. S4**). For the main-chain, clear peaks were observed in the residues from Asn99–Gly102, Ser262**–**Asn263, Phe266, and Tyr277**–**Ala279. Ser262 was hydrogen bonded to Asp187 and Arg189, contributing to its stability. Tyr277**–**Ala279 was located in the C-terminal. Notably, residues Gly100–Tyr104 participate in the formation of the substrate-binding subsites S2–S4 (*62,20*). The substrate-binding subsite S4 is formed between two polypeptide segments of residues Gly100–Tyr104 and Ser132–Tyr137. In contrast, the residues Ser132–Tyr137 showed relatively low r.m.s.d. values. Average *B*-factors in Asn99**–**Tyr104 were also greater than 20 Å^2^, suggesting that the dynamic motion of these residues might help accommodate various types of substrate. On the other hand, the overall crystal structure of the SRX at room temperature (PDB code 4b5l) was highly similar to the SFX structure (**fig. S4**). We observed that the contraction of the cell volume at cryo-temperature was about 4% compared with that from the SFX structure (**Table 1**). The changes in packing will have some effects on the small differences in protein structure.

#### Configuration of hydrogen atoms in active-site regions revealed by SFX

The configurations of the serine protease catalytic triad Ser/His/Asp are important to the catalytic reaction. We have detected, only in the SFX structure, a hydrogen bond with the peaks expected to be hydrogen atoms adjacent to Nε2 of His69 (**Fig. 3**). No peaks were detected near Nε2 of His69 in the SRX structure with similar crystallization conditions at pH 6.5. No significant peak was detected in the σA-weighted m*F*_o_**–**D*F*_c_ map between His69 and Asp39 in the SFX structure, and the distance between Nδ1 of His69 and Oδ1 of Asp39 was 2.70 Å (**Fig. 3b**) and is sufficiently longer than the donor-to-acceptor distance, suggesting proteinase K might not form a low energy barrier hydrogen bond, proposed by the nuclear magnetic resonance (NMR) study (*63*). The carboxylate of Asp39 was charge localized to Oδ1 at the contour level 4.5σ (**Fig. 3b**) and the distance between Cγ and Oδ1 of Asp39 was 1.24 Å, which is shorter than the distance between Cγ and Oδ2 (1.25 Å). These facts suggest that Cγ and Oδ1 are doubly bonded, whereas the Cγ and Oδ2 are singly bonded. These charge distributions were also found in the cryo-cooling SRX data. This suggests that the configurations of active sites formed by Asp39 and His69 are stable against temperature changes. It is difficult to discuss the small structural changes observed in the SFX structure, because there were differences in the experiment temperature, crystallization condition, radiation damage, or a combination of all of these. However, our results indicate that the SFX approach can determine the atomic positions with high accuracy, because crystal irregularities are compensated for by using ~82,000 crystals in different orientations.

In summary, we have obtained the atomic resolution structure of proteinase K at ambient temperature. The resolution of 1.20 Å was attained using ~82,000 indexed patterns from SFX. This atomic resolution structure clearly visualized hydrogen atoms forming hydrogen bond in secondary structures and protonation state of catalytic residues of His69. Electron density distributions from this room temperature SFX provided new structural details such as dynamic motion of substrate binding site and the detailed configurations of active-site regions including water molecules. These directly visualize how enzymes are functioning in physiological conditions. The accurate configurations of atoms acquired by this study will serve to unveil the functional mechanisms of enzymes at ambient temperature.

## Acknowledgments

The XFEL experiments were carried out at the BL3 of SACLA with the approval of the Japan Synchrotron Radiation Research Institute (JASRI) (proposal nos. 2015A8026 and 2015A8048). We performed the synchrotron experiments at the BL26B1 of SPring-8 (proposal no. 2015A1052). **Funding:** This work was supported by the X-ray Free-Electron Laser Priority Strategy Program (MEXT), partly by the Strategic Basic Research Program of Japan Science and Technology Agency (JST), partly by a Grant-in-Aid for Scientific Research from the Japan Society for the Promotion of Science (KAKENHI No. 25650026) and partly by the Platform for Drug Discovery, Informatics, and Structural Life Science (MEXT). C.S. is supported by NRF (2015R1A5A1009962 & 2016R1A2B3010980). The authors thank the SACLA and SPring-8 beamline staff for technical assistance. We thank Allan Nisbet for his useful comments and editing of the manuscript. We are grateful for the computational support from SACLA HPC system and Mini-K super computer system. **Author Contributions:** T.M. and M. Sugahara conceived the research, K.N. and M. Sugahara prepared the microcrystals, T.M. M. Suzuki, S. Inoue, C.S., E.N., R.T., B.M. and M. Sugahara performed the data collection, T.N., T.M. and M. Suzuki performed data processing, T.M. and M. Suzuki solved the structure, T.N. and O.N. developed the data processing pipeline, K.T., Y.J., T.K., T.H. and M.Y. developed the DAPHNIS and detectors, T.M., M. Suzuki, S. Inoue, C.S., T.N., K.N. and M. Sugahara wrote the manuscript with input from all of the coauthors and S. Iwata coordinated the project. **Competing interests:** The authors declare no competing financial interests. **Data and materials availability:** Protein Data Bank: Coordinates and structure factors have been deposited under accession codes 5kxu (SFX) and 5kxv (SRX). Diffraction patterns (SFX) have been deposited to CXIDB under ID #45.

